# Trycycler: consensus long-read assemblies for bacterial genomes

**DOI:** 10.1101/2021.07.04.451066

**Authors:** Ryan R. Wick, Louise M. Judd, Louise T. Cerdeira, Jane Hawkey, Guillaume Méric, Ben Vezina, Kelly L. Wyres, Kathryn E. Holt

**Affiliations:** Department of Infectious Diseases, Central Clinical School, Monash University, Melbourne, VIC, 3004, Australia; Cambridge Baker Systems Genomics Initiative, Baker Heart & Diabetes Institute, Melbourne, VIC, 3004, Australia; Department of Infection Biology, London School of Hygiene & Tropical Medicine, London, WC1E 7HT, UK

## Abstract

Assembly of bacterial genomes from long-read data (generated by Oxford Nanopore or Pacific Biosciences platforms) can often be complete: a single contig for each chromosome or plasmid in the genome. However, even complete bacterial genome assemblies constructed solely from long reads still contain a variety of errors, and different assemblies of the same genome often contain different errors. Here, we present Trycycler, a tool which produces a consensus assembly from multiple input assemblies of the same genome. Benchmarking using both simulated and real sequencing reads showed that Trycycler consensus assemblies contained fewer errors than any of those constructed with a single long-read assembler. Post-assembly polishing with Medaka and Pilon further reduced errors and yielded the most accurate genome assemblies in our study. As Trycycler can require human judgement and manual intervention, its output is not deterministic, and different users can produce different Trycycler assemblies from the same input data. However, we demonstrated that multiple users with minimal training converge on similar assemblies that are consistently more accurate than those produced by automated assembly tools. We therefore recommend Trycycler+Medaka+Pilon as an ideal approach for generating high-quality bacterial reference genomes.

**Data availability:** Supplementary figures, tables and code can be found at: github.com/rrwick/Trycycler-paper

Reads, assemblies and reference sequences can be found at: bridges.monash.edu/articles/dataset/Trycycler_paper_dataset/14890734

## Introduction

Long-read assembly is the process of reconstructing a genome from long sequencing reads (>10 kbp), such as those made by Oxford Nanopore Technologies (ONT) or Pacific Biosciences (PacBio) platforms. ONT’s long-read sequencing platforms are popular for bacterial sequencing due to their low cost per sample^1,2^. Since long reads can span larger genomic repeats than short reads (e.g. reads from Illumina sequencing platforms), long-read assembly can produce larger contigs than short-read assembly^3–6^. For bacterial genomes, it is often possible to produce a long-read-only assembly (an assembly made solely from long-read data) which is complete: one fully assembled contig for each replicon in the genome^7,8^. There are many long-read assemblers appropriate for use on bacterial genomes, including Canu^9^, Flye^10^, Raven^11^ and Redbean^12^. Each has advantages and disadvantages, but in a recent benchmarking study we found Flye to be the best-performing bacterial genome assembler in many metrics^13^.

Since long-read assembly of bacterial genomes can reliably yield chromosome-scale contigs, it is sometimes considered to be a solved problem^14^, with much assembler tool development now focusing on more challenging scenarios such as eukaryotic genomes and metagenomes^15,16^. However, long-read bacterial assemblies are not perfect. Small-scale errors (such as homopolymer-length errors) are commonly discussed and addressed^7,17–19^, but larger-scale errors (tens to hundreds of base pairs) also occur in most assemblies^13^. Even though most bacterial replicons are circular, long-read assemblers often fail to produce cleanly circularised contigs, where the last base in the contig is immediately followed by the first base. Spurious contigs are often present in assemblies (e.g. from contaminant sequences), and small plasmids can be omitted due to their underrepresentation in ONT read sets^20^. Hybrid assembly, which uses both short and long reads, can mitigate some of these problems, but hybrid assemblers also fail to produce error-free genome assemblies^21^, and can introduce confusion if short and long read libraries are not constructed from the same DNA extraction^22^. Long-read assembly of bacterial genomes is therefore not a completely solved problem, and there is still much room for improvement.

As assembly is often the first step in bioinformatic pipelines, assembly errors can have negative implications for downstream analysis. Here we introduce Trycycler, a computational tool which enables high-quality long-read-only assemblies of bacterial genomes. It takes multiple assemblies of the same genome as input and produces a single consensus assembly. Trycycler exploits the fact that while long-read assemblies almost always contain errors, different assemblies of the same genome typically have different errors^13^. Trycycler can therefore combine multiple input assemblies to produce a consensus assembly with fewer errors than any of its inputs.

## Approach and Implementation

The Trycycler pipeline consists of multiple steps which are run separately (overview in **Figure 1**, more detail in **Figure S1**). At the clustering and reconciliation steps, the user may need to make decisions and intervene. This means that Trycycler is not an automated process appropriate for high throughput assembly. Trycycler is implemented in Python and uses the NumPy, SciPy and edlib packages^23–26^.

**Figure 1:**
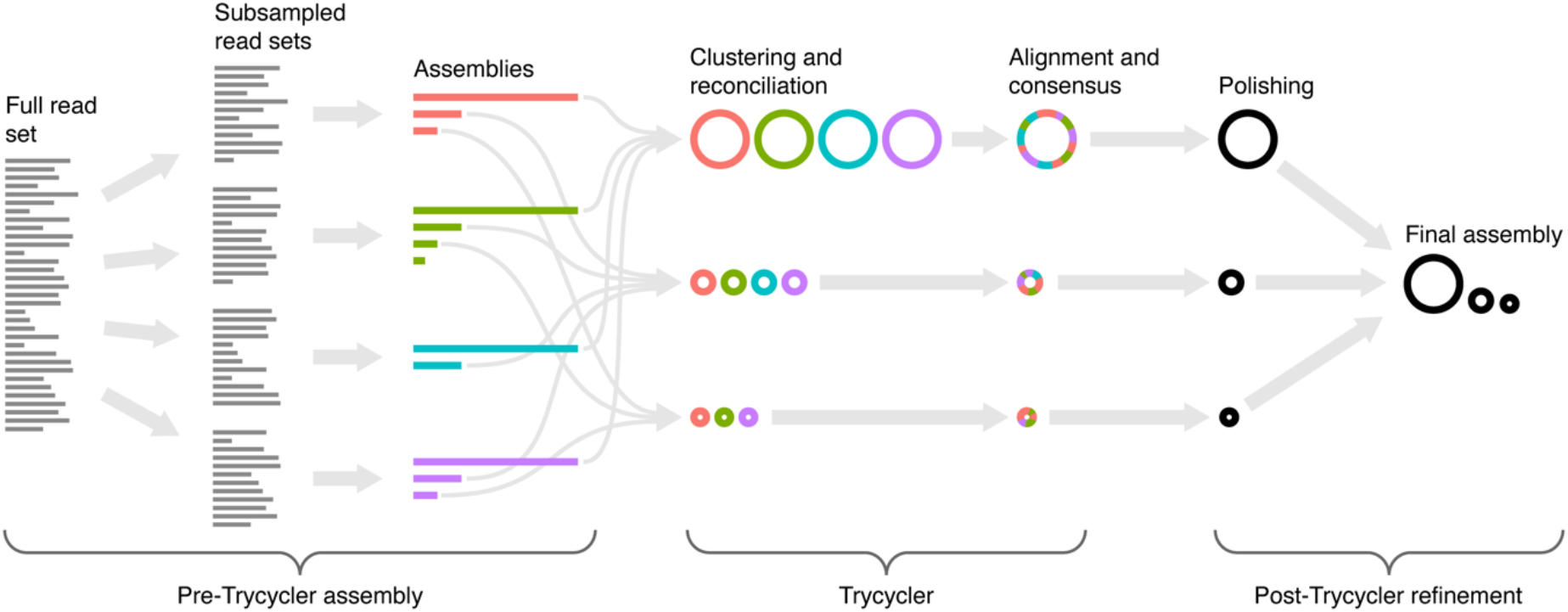
overview of the Trycycler long-read assembly pipeline. Before Trycycler is run, the user must generate multiple completed assemblies of the same genome, e.g. by assembling different subsets of the original long-read set. Trycycler then clusters contigs from different assemblies and produces a consensus contig for each cluster. These consensus contigs can then be polished (e.g. with Medaka) and combined into a final high-quality long-read-only assembly.

Before Trycycler is run, the user must generate multiple input assemblies of the same genome (**Figure S1A**). The input assemblies should be complete: one contig per replicon. If complete assemblies are not possible (e.g. due to insufficient read length) or read depth is shallow (e.g. <25× depth), then Trycycler is not appropriate. We recommend users generate 12 independent input assemblies, but this value can be adjusted down (to save computational time) or up (to improve robustness). It is desirable to maximise the independence of the input assemblies, as this will reduce the chance that the same error will occur in multiple assemblies. One way to achieve such independence is to use multiple assemblers, as different assembly algorithms can lead to different assembly errors^13^. For example, in the tests reported here we used Flye^10^, Miniasm/Minipolish^13^, Raven^11^ and Redbean^12^. Random read subsampling can provide further independence, where each assembly is generated from a different subsample of the full read set (Trycycler v0.5.0 has a ‘subsample’ command to facilitate this). Deeper long-read sets are therefore desirable, as they enable more independent subsets.

The first step in the Trycycler pipeline is contig clustering (**Figure S1B**). It aims to group contigs of the same replicon from different input assemblies, so subsequent steps can be carried out on a per-replicon basis. For example, if the genome in question had one chromosome and one plasmid, then Trycycler clustering should produce two clusters: one for the chromosomal contigs and one for the plasmid contigs. To make clusters, Trycycler conducts complete-linkage hierarchical clustering on all pairwise Mash distances between contigs^27^. To aid interpretation, a FastME tree is built using the pairwise distances^28^. After clustering is complete, the user must decide which clusters are valid (i.e. represent completely assembled replicons in the genome) and which are invalid (i.e. represent incomplete, misassembled or spurious sequences) – a key point of human judgement in the Trycycler process.

The next step is to ‘reconcile’ each cluster’s contig sequences with each other (**Figure S1C**). This involves converting sequences to their reverse complement as necessary to ensure that all sequences in the cluster are in the same orientation. Most bacterial replicons are circular, so Trycycler aligns the start and end of each contig to the other contigs in the cluster to determine if bases need to be added or removed for clean circularisation (can be disabled for linear replicons by using the --linear option). It then rotates each sequence to begin at the same position. Some gene sequences (e.g. *dnaA* and *repA*) are often used as starting positions in complete genomes, so Trycycler contains a database of these genes and will preferentially use them as the contig starting position (see **Methods**). If no sequence from this database is found (with ≥95% coverage and ≥95% identity), Trycycler will use a randomly chosen unique sequence instead.

After reconciliation, each cluster’s sequences will have a consistent strand and starting position, making them appropriate for global multiple sequence alignment (**Figure S1D**). To improve computational performance, Trycycler partitions the sequences into smaller pieces, using 1 kbp pieces with each piece extended as necessary to ensure that the boundaries between pieces do not start/end in repetitive regions. It uses MUSCLE^29^ to produce a multiple sequence alignment for each piece, and then stitches the pieces together to produce a single multiple sequence alignment for the full cluster sequences. Trycycler then aligns the entire read set to each contig sequence so it can be assigned to a particular cluster (**Figure S1E**).

The final step in Trycycler’s pipeline is the generation of a consensus sequence for each cluster (**Figure S1F**). It does this by dividing the multiple sequence alignment into regions where there is or is not any variation. For all regions where there is variation, Trycycler must choose which variant will go into the consensus. The best variant is defined as the one with the minimum total Hamming distance to the other variants, an approach which favours more common variants. In the event of a tie between two variants, Trycycler aligns the cluster’s reads to each possibility and chooses the one which produces the largest total alignment score – i.e. the variant which is in best agreement with the reads. The final Trycycler consensus sequence for the cluster is produced by taking the best variant for each region of variation in the multiple sequence alignment.

After Trycycler finishes, it is recommended to perform long-read polishing on its consensus sequences (**Figure S1G**). Polishing is not incorporated into Trycycler, as that step can be specific to the long-read sequencing technologies used, e.g. Medaka^30^ polishing for ONT assemblies. If short reads are available, short-read polishing (e.g. with Pilon^31^) can also be performed to further improve assembly accuracy.

The code and documentation for Trycycler v0.3.3 (the version used to generate the assemblies in this manuscript) are available at the DOI 10.5281/zenodo.3966493. The current version of Trycycler (v0.5.0) is available on GitHub (github.com/rrwick/Trycycler).

## Results

### Performance on simulated reads

*In silico* read simulation allows for a straightforward test of assembly accuracy against a ground truth: reads are generated from a reference genome, the reads are assembled, and the resulting assembly is compared back to the original reference sequence. For this analysis, we simulated short and long reads from 10 reference genomes which belong to the 10 most common bacterial species in RefSeq (**Table S1**). We assembled each genome with long-read-only approaches (Miniasm/Minipolish^13^, Raven^11^, Flye^10^ and Trycycler), long-read-first hybrid approaches (Pilon^31^ polishing of each long-read-only assembly) and a short-read-first hybrid approach (Unicycler^21^). We quantified the accuracy of each assembly’s chromosomal contig using two main metrics: mean identity and worst-100-bp identity (the minimum identity observed among 100-bp sliding windows).

Comparing only the long-read assemblers to each other (Flye, Miniasm/Minipolish and Raven), it was clear that Flye performed best (**Figure S2**). This was true both before Pilon polishing with short reads (mean identity Q41 vs Q38; mean worst-100-bp-identity 95.8% vs 50.8–90.9%) and after Pilon polishing (mean identity Q57 vs Q42–Q55; mean worst-100-bp identity 96.1% vs 50.8–95.7%). Our main results therefore exclude Miniasm/Minipolish and Raven, leaving only the best-performing long-read assembler: Flye.

**Figure 2** shows the mean assembly identities and worst-100-bp assembly identities from each approach, using 10 simulated read sets. In both metrics, Trycycler reliably produced higher-quality assemblies than Flye (mean identity Q51 vs Q41; mean worst-100-bp identity 99.5% vs 95.8%). This result also held true for long-read-first hybrid assemblies, where Trycycler+Pilon outperformed Flye+Pilon (mean identity Q74 vs Q57; mean worst-100-bp identity 99.9% vs 96.1%). Unicycler’s short-read-first hybrid assemblies performed notably worse than the long-read-first hybrid approaches (mean identity Q25; mean worst-100-bp identity 76.5%).

**Figure 2:**
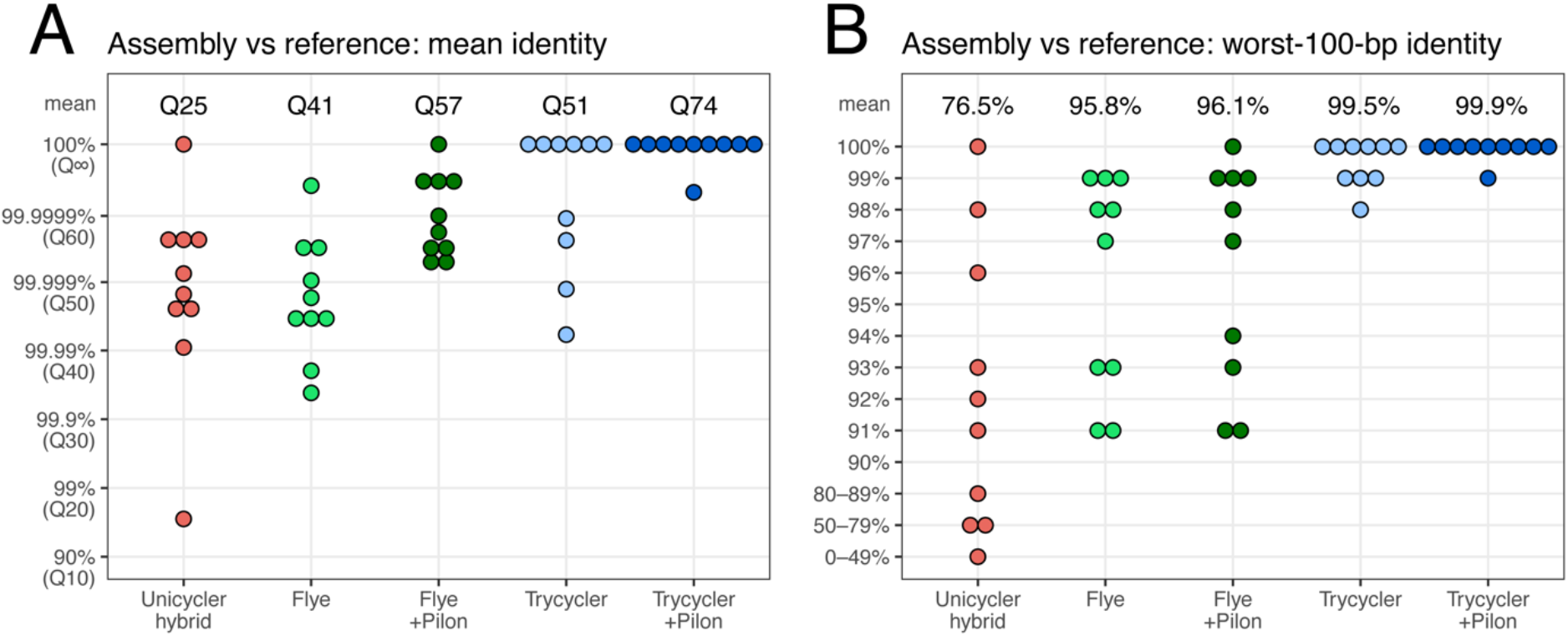
results for the tests using simulated reads. For 10 reference genome sequences, we simulated both short and long reads. The read sets were then assembled with Unicycler (short-read-first hybrid assembly), Flye (long-read-only assembly), Flye+Pilon (long-read-first hybrid assembly), Trycycler (long-read-only assembly) and Trycycler+Pilon (long-read-first hybrid assembly). Each assembled chromosome was aligned back to the reference chromosome to determine the mean assembly identity (A) and the worst identity in 100-bp sliding windows (B). For long-read-only assembly, Trycycler consistently achieved higher accuracy than Flye. Trycycler+Pilon (i.e. using Pilon to polish the Trycycler assembly with short reads) achieved the highest accuracy and did better than alternative hybrid approaches (Unicycler and Flye+Pilon).

### Performance on real reads

Since simulated reads cannot perfectly emulate real sequencing^32^, we also tested assembly methods with real read sets. We chose seven bacterial isolates for this study (**Table S2**), each belonging to a different bacterial species with clinical relevance. The challenge with real reads is the absence of a clear ground truth against which to compare assemblies. To circumvent this issue, we instead produced two independent sets of long+short (ONT+Illumina) reads for each test organism. In brief, a single DNA extraction from each organism was used to prepare two ONT libraries (one ligation, one rapid), and a single Illumina library (the results of which were divided into two non-overlapping read sets); full details are described in **Methods**. For each assembly method, we compared the assembly from read set A to the assembly of read set B, differences between them indicating assembly errors. While this approach could suffer from false negatives if both assemblies contained the same error, it cannot suffer from false positives, as wherever two assemblies of the same genome differ, at least one of the two must be in error.

We tested the same assemblers as were used in the simulated-read tests but added an additional long-read polishing step with Medaka, an ONT-specific polishing tool. We therefore produced unpolished long-read-only assemblies (with Miniasm/Minipolish, Raven, Flye and Trycycler), polished long-read-only assemblies (the same assemblers plus Medaka), long-read-first hybrid assemblies (the same assemblers plus Medaka and short-read polishing with Pilon) and short-read-first hybrid assemblies (with Unicycler). Each assembly approach was used on both read set A and read set B for each of the test organisms.

Assembly accuracy was quantified using the metrics from the simulated-read tests: mean identity and worst-100-bp identity. Instead of being based on an assembly-to-reference alignment (as was done for the simulated-read tests), these metrics used an alignment of the read-set-A-assembled chromosome to the read-set-B-assembled chromosome. For the *Serratia marcescens* genome, read set B failed to produce a complete chromosome with most assembly methods (due to long genomic repeats and a short read N50, see **Table S2**), so this genome was excluded, leaving six genomes in the analyses. As was the case for the simulated-read tests, Flye assemblies were higher quality than Miniasm/Minipolish and Raven assemblies at all polishing stages (**Figure S3**): unpolished (mean identity Q34 vs Q28–Q32; mean worst-100-bp identity 81.8% vs 20.2–21.8%), Medaka-polished (mean identity Q40 vs Q30–Q35; mean worst-100-bp identity 94.7% vs 28.2–38.0%) and Medaka+Pilon-polished (mean identity Q56 vs Q31–Q37; mean worst-100-bp identity 94.7% vs 28.2–40.0%). Flye was also the only long-read assembler to produce completed chromosomes for both read sets of all six genomes, so Miniasm/Minipolish and Raven were excluded from our main results.

Since the mean identity and worst-100-bp identity metrics could fail to identify all assembly errors in the real-read tests, we also used two other approaches for assessing the quality of *de novo* assemblies. The first was ALE^33^, which uses short-read alignments to the assembled sequence to produce a likelihood score for that assembly (higher scores being better), which we normalised for each genome to produce a z-score. Mapping accuracy, evenness of read depth and evenness of insert size extracted from the short-read alignments are all used by ALE to generate likelihood scores. The second *de novo* assessment approach was IDEEL^34,35^, which compares the length of predicted proteins in the assembly to a database of known proteins. Indel errors in the assembly cause frameshifts in coding sequences leading to truncations, so an error-prone assembly will tend to have predicted proteins which are shorter than their best-matching known proteins. We quantified the fraction of predicted proteins in each assembly which were ≥95% the length of their best-matching known protein (higher fractions being better).

**Figure 3** shows the real-read results: mean identity, worst-100-bp identity, ALE z-scores and IDEEL full-length proteins. In the mean identity metric, Trycycler performed better than Flye at all levels of polishing (Q37 vs Q34 before polishing; Q42 vs Q40 after Medaka polishing; Q62 vs Q56 after Medaka+Pilon polishing). This advantage was also apparent in the worst-100-bp identity metric (96.7% vs 81.8% before polishing; 97.0% vs 94.7% after Medaka polishing; 98.3% vs 94.7% after Medaka+Pilon polishing). Both long-read-first hybrid approaches (Flye+Medaka+Pilon and Trycycler+Medaka+Pilon) outperformed Unicycler’s short-read-first hybrid assemblies (mean identity Q34 and worst-100-bp identity 23.5%). The ALE results are consistent with the identity metrics: Trycycler assemblies had higher mean ALE z-scores than Flye assemblies at all polishing levels (–1.031 vs –1.873 before polishing; 0.419 vs 0.235 after Medaka polishing; 0.828 vs 0.806 after Medaka+Pilon polishing) and long-read-first hybrid assemblies were superior to Unicycler assemblies (mean ALE z-score of 0.617). IDEEL results showed the same trend, with Trycycler assemblies having more full-length proteins than Flye assemblies (78.3% vs 72.3% before polishing; 93.8% vs 91.8% after Medaka polishing), but all hybrid assemblies performed equivalently in this metric (97.6%).

**Figure 3:**
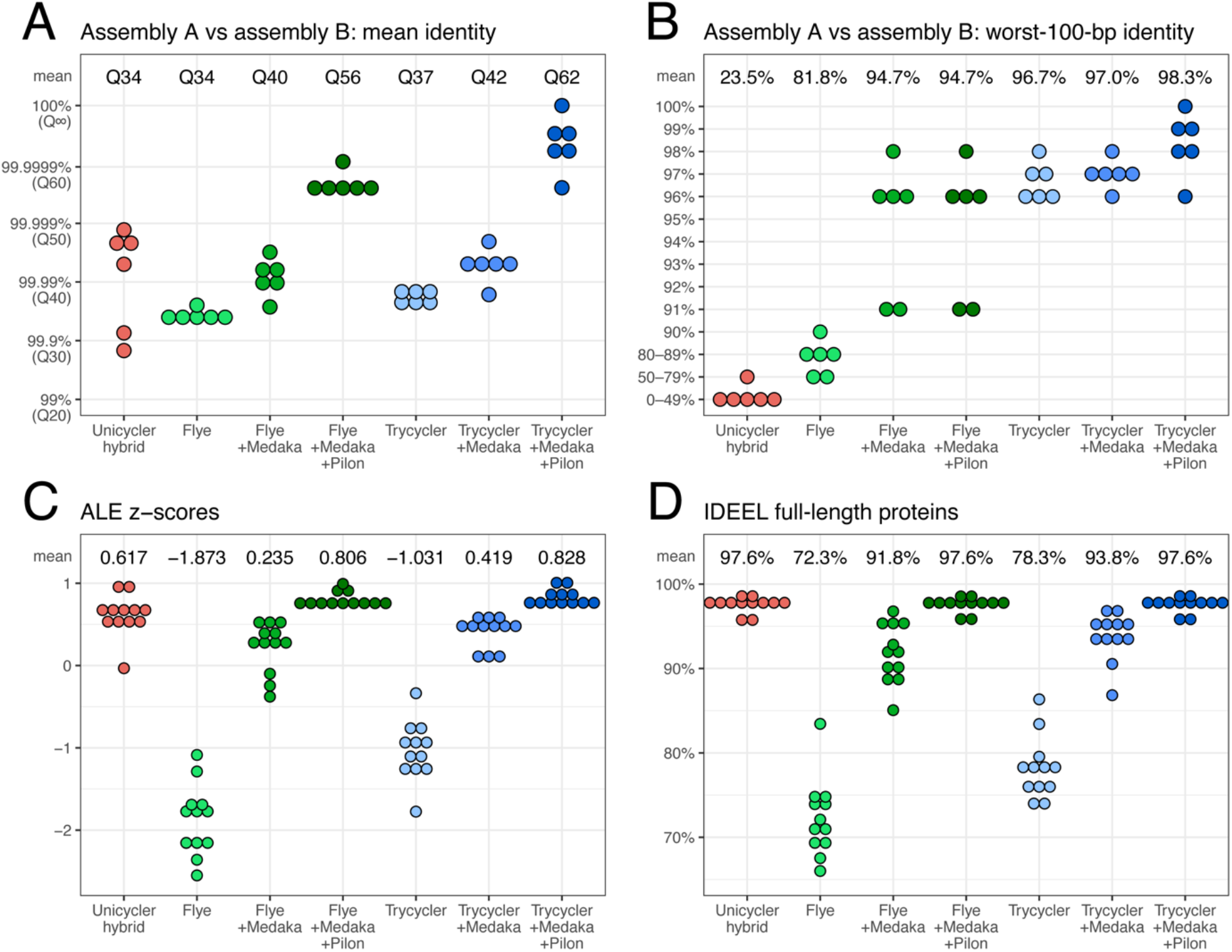
results for the real-read tests. For six genomes, we produced two independent hybrid read sets from the same DNA extraction. The read sets were then assembled with Unicycler (short-read-first hybrid assembly), Flye (long-read-only assembly), Flye+Medaka (long-read-only assembly), Flye+Medaka+Pilon (long-read-first hybrid assembly), Trycycler (long-read-only assembly), Trycycler+Medaka (long-read-only assembly) and Trycycler+Pilon (long-read-first hybrid assembly). For each genome and each assembly approach, we aligned the two independently assembled chromosomes to each other to determine the mean assembly identity (A) and the worst identity in 100-bp sliding windows (B). For long-read-only assembly, Trycycler consistently achieved higher accuracy than Flye (both before and after Medaka polishing). Trycycler+Medaka+Pilon achieved the highest accuracy and did better than alternative hybrid approaches (Unicycler and Flye+Medaka+Pilon). We also assessed the accuracy of each of the 12 assembled chromosomes using ALE (C) and IDEEL (D). ALE assigns a likelihood score (transformed into z-scores on a per-genome basis) to each assembly based on its concordance with the Illumina read set. IDEEL identifies the proportion of predicted proteins which are ≥95% the length of their best-matching known protein in a database.

### Type and location of errors

**Figure S4** shows the positions of errors in the assemblies of each of the 16 genomes (10 simulated and six real), with repetitive regions of the genomes indicated. Errors in long-read-only assemblies (Flye, Flye+Medaka, Trycycler and Trycycler+Medaka) were distributed across the genomes, occurring in both repeat and non-repeat sequences. Long-read-first hybrid assemblies (Flye+Medaka+Pilon and Trycycler+Medaka+Pilon) usually had higher error rates in repeat sequences, and in many cases, there were no errors in the non-repeat sequences of the genome. Short-read-first hybrid assemblies (Unicycler) often had clusters of errors which occurred in both repeat and non-repeat sequences. Indel errors were more common than substitution errors for all assemblers: 44% of total errors were insertions, 47% were deletions and 9% were substitutions. For the real reads, Flye assemblies sometimes had local spikes in error rates (indicating a more serious error or a cluster of errors) before Medaka polishing, but these spikes were not present after Medaka polishing. Trycycler assemblies did not suffer from this same problem. Flye assemblies often had errors at the position corresponding to the original start/end of the contig.

The Flye errors at the start/end of the contig were caused by imperfect circularisation: missing or duplicated bases at the start/end of a circular contig, a phenomenon we described in greater detail in a previous benchmarking study of long-read assemblers^13^. These errors were not corrected by Medaka or Pilon because those tools are not aware of contig circularity, i.e. that the contig’s last base should immediately precede its first base. Since our analysis involved normalising all assemblies to a consistent starting position (required for global alignment), missing/duplicated bases at the start/end of a contig registered as a middle-of-the-sequence indel error in our tests. These indel errors reduced the mean identity and, if large enough, the worst-100-bp identity as well.

To assess the effect of circularisation errors on Flye accuracy, we manually fixed the circularisation of all Flye assemblies using the original reference sequence (in the simulated-read tests) or the Trycycler+Medaka+Pilon assembly (in the real-read tests). Of the 22 Flye assemblies (10 from simulated reads, 12 from real reads), four had perfect circularisation, five had duplicated bases and 13 had missing bases. The worst Flye circularisation error was a 13 bp deletion, and the mean magnitude of Flye circularisation errors was 3.7 bp (**Tables S1 and S2**). We then reran our analyses using the fixed-circularisation version of Flye assemblies, and the results are shown in **Figure S5** for simulated reads and **Figure S6** for real reads. Flye performed better in these results, especially in the worst-100-bp identity metric, indicating that in many cases, the circularisation error was the largest single error in the Flye assembly. However, Trycycler still produced more accurate assemblies than Flye at each polishing stage (unpolished, Medaka-polished and Pilon-polished).

### Consistency of Trycycler results

Trycycler is not a fully automated pipeline – it requires human judgement and intervention. This raised the question of how well it performs in the hands of different users. To answer this, we recruited five researchers who were experienced in bioinformatics but not involved in Trycycler development. They were given an ONT read set for each of the six genomes used in the real-read tests and tasked with producing a Trycycler assembly without any assistance from the Trycycler developer (using only the Trycycler documentation to guide them). We then compared the resulting assemblies, looking at both presence/absence of contigs as well as chromosomal sequence identity (**Figure 4**).

**Figure 4:**
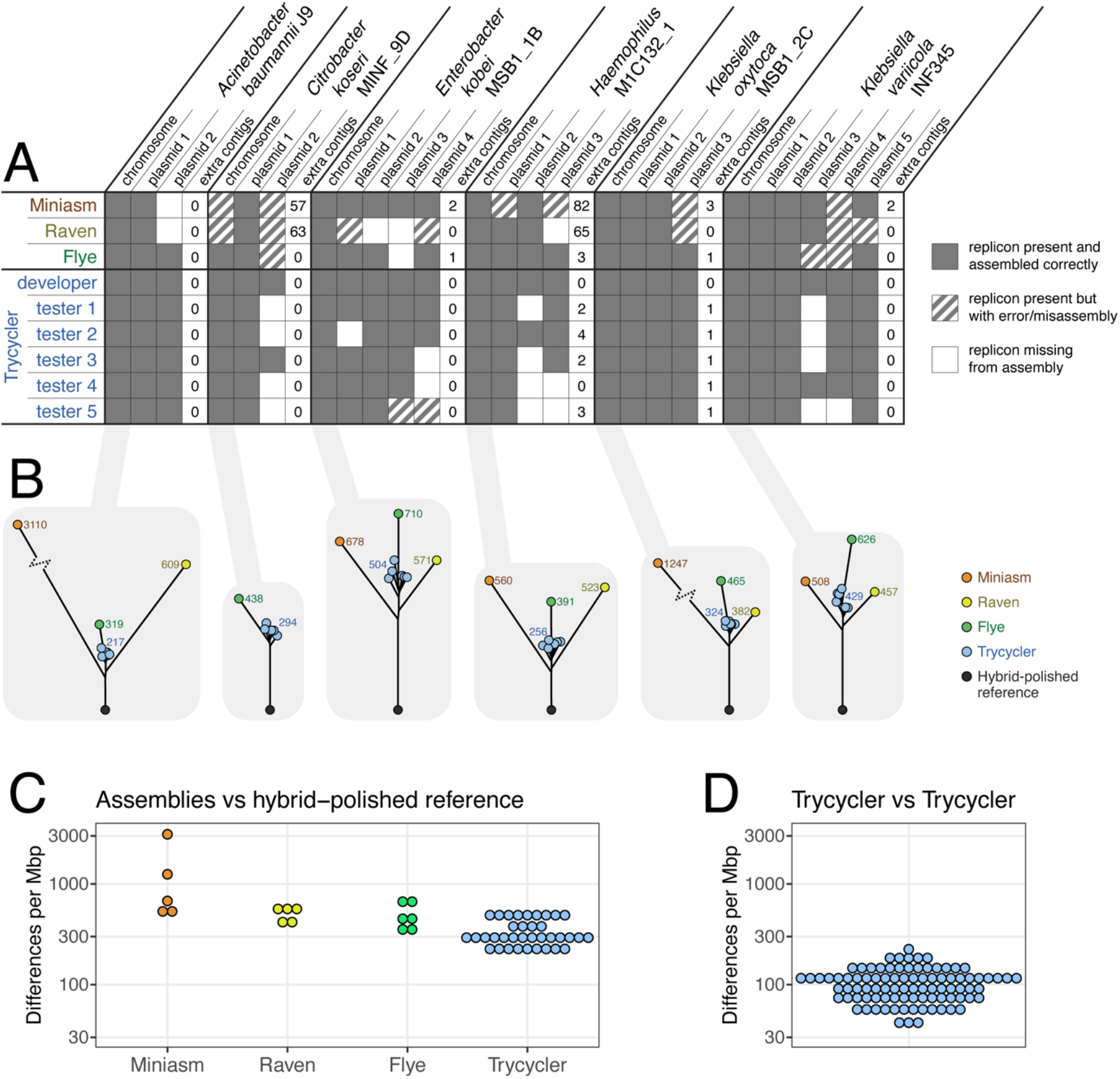
results for the multi-user test which assessed the consistency of Trycycler assemblies when run by different users. Results include assemblies from three different long-read assemblers (Miniasm/Minipolish, Raven and Flye, all automated and deterministic for a given set of reads and parameters, i.e. independent of user) and Trycycler assemblies from six different users (the developer of Trycycler and five testers). A: Presence/absence matrix for the replicons in the test genomes. Each replicon was classified as either present in the assembly, absent from the assembly, or present but with an error/misassembly (see Table S3 for more detail). The number of additional contigs (e.g. spurious or contaminant sequences) is also indicated for each assembly. All Trycycler assemblies contained an accurate chromosome, and only one Trycycler assembly contained misassemblies. However, in many cases the Trycycler testers excluded a true plasmid (most commonly a small plasmid) or included an additional plasmid (most commonly constructed from cross-barcode contaminating reads). B: Neighbour-joining trees of all available assemblies for each of the chromosomes, based on pairwise alignment distances. Hybrid-polished (Medaka+Pilon) versions of the developer’s Trycycler assemblies were included as reference sequences. The values indicate the number of single-bp differences per Mbp between each assembly and the polished reference (values for Trycycler are the mean of all six Trycycler assemblies). For each genome, the Trycycler assemblies cluster tightly and are closer to the polished reference than those from other long-read assemblers. C: Differences between each assembled chromosome and the hybrid-polished reference. Values are single-bp differences per Mbp of sequence. Trycycler assemblies contain fewer differences, on average, compared to the single-assembler assemblies. D: Pairwise differences between Trycycler assemblies of each chromosome. Values are single-bp differences per Mbp of sequence, and there are 90 values (6 genomes × 15 unique pairwise combinations per genome).

The main source of variation between different users’ Trycycler assemblies was the inclusion/exclusion of plasmid contigs (**Figure 4A**). Small plasmids often pose problems for long-read assemblers, and this caused them to sometimes be excluded by Trycycler users. Contaminant plasmid contigs (e.g. cross-barcode contamination) were sometimes included in Trycycler assemblies. Replicons with a large-scale error or misassembly occurred in many of the single-assembler assemblies (from Miniasm/Minipolish, Raven and Flye). These errors included fragmented replicons (e.g. splitting one replicon sequence between two contigs), doubling a replicon in a single contig (e.g. assembling a 6 kbp plasmid into a 12 kbp contig), large-scale circularisation problems (e.g. 80 kbp of start/end overlap), and redundant contigs (e.g. producing five contigs for a single replicon). This type of error was very rare in the Trycycler assemblies (present in only one case). Detailed descriptions of all such errors are in **Table S3**.

To assess the consistency of assembled sequences, we built a neighbour-joining tree (based on pairwise alignment distances) of the assembled chromosomes for each of the six genomes (**Figure 4B**). The developer’s Trycycler+Medaka+Pilon assembly was included as a reference sequence, as the real-read test results (**Figure 3**) indicate these to be the most accurate representation of the genomes. For each test isolate, the Trycycler assemblies generated by different users were closer to the reference sequence than any of the (automated) single-assembler assemblies (**Figure 4C**), and there were comparatively few differences between Trycycler assemblies from different users (**Figure 4D**). All differences between Trycycler assemblies generated by different users were small-scale: most were only single-bp differences, and the largest difference was a 4-bp indel in a tandem repeat (**Table S3**). The most common difference was a 1-bp discrepancy in the length of a homopolymer sequence (accounted for 78.5% of all Trycycler- vs-Trycycler sequence differences).

## Discussion

By combining multiple input assemblies into a consensus sequence, Trycycler produced the most accurate long-read-only assemblies in our study (**Figures 2–4**). Trycycler assemblies only contained small-scale errors (i.e. their accuracy in a 100-bp sliding window remained high), while assemblies produced by single assemblers often contained medium-to-large scale errors (**Figures 2–3**). Trycycler also helped to guard against inexact circularisation, inclusion of spurious contigs and exclusion of genuine contigs. However, Trycycler requires deeper long-read sets (to allow for multiple independent input assemblies via read subsampling), more computational resources and more human input than single-assembler assemblies.

Creating a Trycycler assembly often requires human judgement and manual intervention, particularly after Trycycler’s clustering step where users must decide which contig clusters are valid (represent true replicons in the genome) and which are invalid (spurious, misassembled or contaminant sequences). Our multi-user consistency test showed that this step was a significant source of variability in Trycycler results, manifesting as missing/extra replicons in the assembly, a problem exacerbated by cross-barcode contamination and the fact that long-read assemblers often struggle with small plasmid sequences. This demonstrates that user skill and experience is an important factor in producing an ideal Trycycler assembly. To mitigate this concern, we have provided extensive documentation for Trycycler, with sample data, example analyses and FAQs to guide users. Notably though, Trycycler chromosome sequences generated by different users were more similar to one another than to any of the sequences generated by the deterministic single assemblers (**Figure 4B**).

Polishing is a post-assembly processing step which aims to improve sequence accuracy, and it can be carried out using either long or short reads. Our study showed that Medaka, a long-read polishing tool for ONT reads, was able to fix approximately half of the errors in long-read assemblies. Medaka was also effective at repairing many of the worst errors in a Flye assembly, making Flye+Medaka assemblies nearly as accurate as Trycycler+Medaka assemblies. Subsequent short-read polishing with Pilon was able to bring sequence identity close to 100%, with most of the remaining unfixed errors residing in genomic repeats (where short-read alignment is unreliable). Our study also found short-read-first hybrid assembly (short-read assembly followed by long-read scaffolding, as performed by Unicycler) to be less reliable than long-read-first hybrid assembly (long-read assembly followed by short-read polishing). However, in cases where short reads are deep but long reads are shallow (not tested in this study), Unicycler is likely to perform better, as this was the case it was designed for^21^.

The goal of any assembly approach is to produce a representation of the underlying genome with the fewest errors. Assuming there is a single, unambiguous underlying genome (i.e. no genomic heterogeneity), the ideal result is a base-for-base exact match of the genome: a perfect assembly. Our study shows that for bacterial genomes, a Trycycler+Medaka+Pilon approach can deliver assemblies which are very close to this goal: approximately one error per 2 Mbp, equivalent to two errors in an *E. coli* genome. Future improvements in sequencing technologies, basecalling and assembly/polishing algorithms may make perfect bacterial assemblies a reality, and only when this is reliably achievable can we truly call bacterial genome assembly a ‘solved problem’.

## Methods

### Starting gene database

To generate Trycycler’s database of preferred contig-starting gene sequences, we produced consensus sequences of common genes at the start of completed contigs on RefSeq. All completed bacterial genomes on RefSeq were downloaded, and the name of the first gene in each contig was extracted. These names were tallied and sorted to produce a list of common starting gene names, e.g. ‘Chromosomal replication initiator protein DnaA’ and ‘Replication initiation protein’. The gene sequences with these names were extracted and clustered using complete-linkage hierarchical clustering (coverage threshold of 100% and sequence identity threshold of 95%). We then produced an ancestral state reconstruction consensus sequence for each cluster using MUSCLE^29^, FastTree^36^ and TreeTime^37^ to generate the final set of 7171 contig starting sequences.

### Simulated-read tests

One reference genome was used from each of the 10 most common bacterial species in RefSeq: *Escherichia coli*, *Salmonella enterica*, *Staphylococcus aureus, Streptococcus pneumoniae*, *Klebsiella pneumoniae*, *Mycobacterium tuberculosis*, *Pseudomonas aeruginosa*, *Listeria monocytogenes*, *Neisseria meningitidis* and *Campylobacter jejuni* (**Table S1**). Badread (v0.1.5) was used to simulate a long-read set for each genome^32^. The parameters (read length, read accuracy, chimera rate, etc.) were varied between sets to test a variety of inputs. To ensure assemblability, all read sets were 100× depth or greater and the mean read length was longer than the longest repeat in the genome (as determined by a self-vs-self MUMmer alignment^38^). To simulate short reads for each genome, we used ART (v2016-06-05) and the parameters (simulation profile, depth, read length and fragment length) were varied between genomes^39^. Simulation parameters and summary statistics for each simulated read set are in **Table S1**. Before assembly, we conducted quality-control filtering using fastp v0.20.1^40^ for short reads (using default parameters) and Filtlong^41^ v0.2.0 for long reads (using a minimum read length of 1 kbp and a kept-base percentage of 95%). Simulated reads are available in **Supplementary data**.

Unicycler^21^ (v0.4.8) assemblies were conducted on each hybrid (short and long) read set using the --no_correct option to disable read error correction because the documentation for SPAdes^42^ (Unicycler’s underlying assembler) recommends disabling read error correction for high-depth whole genome bacterial reads. Miniasm/Minipolish^4^ (v0.3/v0.1.3), Raven^43^ (v1.2.2) and Flye^10^ (v2.7.1) assemblies were conducted on each long-read set using default parameters for each. Trycycler assemblies were performed using default parameters and following the procedure outlined in the Trycycler documentation (12 input assemblies made from subsampled read sets of 50× depth). Versions of Flye assemblies with repaired start/end indels were produced by manually comparing the Flye sequence to the reference genome sequence. All long-read-only assemblies were then polished with Bowtie2^44^ (v2.3.4.1) and Pilon^31^ (v1.23). For Bowtie2 read alignment, we set min/max fragment lengths using values from the Unicycler assembly log (1^st^ and 99^th^ fragment size percentiles). We conducted multiple rounds of Bowtie2+Pilon polishing, stopping when it ceased to make any changes or at five rounds, whichever came first. See **Supplementary data** for the exact assembly and polishing commands used. Complete chromosomal assembly was assessed by a manual inspection of the assembly graphs and looking for an appropriately sized circular contig. All simulated read assemblies produced a single chromosomal contig with one exception: the Unicycler assembly for the *N. meningitidis* genome. However, the Unicycler assembly graph for *N. meningitidis* contained a single unbranching loop, so we merged the resulting contigs to produce a single chromosomal sequence.

To quantify the accuracy of the assemblies, we manually extracted the chromosomal contig from each assembly’s graph. We then made the contig consistent with the reference sequence by normalising the strand (changing the sequence to its reverse complement if necessary) and starting position (moving bases from the beginning of the contig to the end) to match the reference genome. The pairwise_align.py script (available in **Supplementary data**) was then used to perform a global sequence alignment between each contig and its reference sequence using the edlib library^26^. From this alignment, we produced two metrics: the mean sequence identity (the number of matching bases divided by the full alignment length) and the worst-100-bp identity (the minimum number of matching bases in a 100-bp sliding window over the alignment). We then used the error_positions.py script (available in **Supplementary data**) to identify the position, type and size of each assembly error, and quantify the accuracy in repeat and non-repeat sequences.

### Real-read tests

The seven bacterial isolates used in this study each belong to a different species: *Acinetobacter baumannii*, *Citrobacter koseri*, *Enterobacter kobei*, an unnamed *Haemophilus* species (given the placeholder name *Haemophilus sp002998595* in GTDB R202^45,46^), *Klebsiella oxytoca*, *Klebsiella variicola* and *Serratia marcescens*. Isolates were cultured overnight at 37°C in Luria-Bertani broth and DNA was extracted using GenFind v3 according to the manufacturer’s instructions (Beckman Coulter). The same DNA extract was used to sequence each isolate using three different approaches: ONT ligation, ONT rapid and Illumina. For ONT ligation, we followed the protocol for the SQK-LSK109 ligation sequencing kit and EXP-NBD104 native barcoding expansion (Oxford Nanopore Technologies). For ONT rapid, we followed the protocol for the SQK-RBK004 rapid barcoding kit (Oxford Nanopore Technologies). All ONT libraries were sequenced on MinION R9.4.1 flow cells. For Illumina, we followed a modified Illumina DNA Prep protocol (catalogue number 20018705), whereby the reaction volumes were quartered to conserve reagents. Illumina libraries were sequenced on the NovaSeq 6000 using SP reagent kits v1.0 (300 cycles, Illumina Inc.), producing 150 bp paired-end reads with a mean insert size of 331 bp. All ONT read sets were basecalled and demultiplexed using Guppy v3.6.1. The resulting Illumina read pairs were shuffled and evenly split into two separate read sets. We then produced two non-overlapping hybrid read sets (A and B) for each genome. Read set A consisted of the ONT ligation reads plus half of the Illumina reads. Read set B consisted of the ONT rapid reads plus the other half of the Illumina reads. All reads are available in **Supplementary data**.

Read sets A and B for each isolate (14 total read sets) were subjected to the same read QC and assembly methods as were used for the simulated read sets, to generate long-read-only and hybrid assemblies for comparison. Versions of Flye assemblies with repaired start/end indels were produced by manually comparing the Flye sequence to the Trycycler+Medaka+Pilon assembly. We separately polished each contig from each long-read-only assembly using Medaka, using Trycycler-partitioned reads, the r941_min_high_g360 model (to match the basecalling model used) and default parameters. We then polished each long-read Medaka-polished assembly using Pilon as described above. See **Supplementary data** for the exact assembly and polishing commands used.

To quantify the accuracy of the resulting assemblies, we manually extracted the chromosomal contig from each, where possible. The *Serratia marcescens* 17-147-1671 read set B assemblies usually failed to produce a complete chromosomal contig (only Unicycler succeeded), so that genome was excluded from further analyses. For the six remaining genomes, we normalised all chromosomes to the same strand and starting position, then used the pairwise_align.py script (available in **Supplementary data**) to perform a global sequence alignment between read set A and read set B chromosomes using the edlib library^26^. From this alignment, we produced the same metrics as were used in the simulated-read tests: mean sequence identity and worst-100-bp identity. We then used the error_positions.py script (available in **Supplementary data**) to identify the position, type and size of each assembly error, and quantify the accuracy in repeat and non-repeat sequences.

To produce ALE scores, we aligned the full short-read set (i.e. before it was split into read sets A and B) to each assembled chromosome using Bowtie2^44^ (v2.3.4.1). The alignments were then given to ALE to produce a single likelihood score^33^. ALE analyses were done on each assembly, so generated 12 values (read sets A and B for each of the six genomes) for each assembly method. ALE scores are not an absolute metric of assembly quality, only a relative metric for comparing different assemblies of the same genome. We therefore normalised the ALE scores using the mean and standard deviation for each genome to produce ALE z-scores.

IDEEL analysis of genomes requires a protein database, so we download all UniProt/TrEMBL^47^ release 2020_05 sequences. From this we built a Diamond^48^ (v2.0.4) index which was used by IDEEL^34^. Predicted proteins in the assembly were classified as full-length if IDEEL found them to be ≥95% the length of the best-matching known protein in the database. IDEEL analyses were done on each assembly and generated 12 values (read sets A and B for each of the six genomes) for each assembly method.

### Multi-user consistency tests

Each of the five Trycycler testers were given the ONT rapid read set for the six genomes used in the real-read tests (all real genomes excluding *Serratia marcescens* 17-147-1671) and produced one Trycycler assembly (without Medaka or Pilon polishing) for each. The number of input assemblies and which assemblers were used are available in **Table S3**. We then compared the assemblies produced by single tools (Flye, Raven and Miniasm/Minipolish), by Trycycler (from the developer and the five testers), and a hybrid-assembled reference (the developer’s Trycycler+Medaka+Pilon assembly).

For each genome, we clustered the contigs from all assemblies (using Trycycler cluster), and using the developer’s Trycycler assembly as the reference, we classified the genome replicons for each assembly as either present, present with misassemblies or absent (**Table S3**). Each chromosome was rotated to a consistent starting position and a multiple sequence alignment was performed (using Trycycler MSA). We then extracted pairwise distances from the alignment (using the msa_to_distance_matrix.py script, available in **Supplementary data**) and built a FastME^28^ tree from the distances. The distances were then normalised to the genome size (using the normalise_distance_matrix_to_mbp.py script, available in **Supplementary data**) to quantify the differences between each assembled chromosome for each of the genomes.

